# Multiple Mechanisms of HIV-1 resistance to PGT135 in Chinese Subtype B’ Slow Progressor

**DOI:** 10.1101/2020.12.22.424087

**Authors:** Shasha Sun, Sen Zou, Yuanyuan Hu, Ying Liu, Li Ren, Yanling Hao, Xintao Hu, Yuhua Ruan, Liying Ma, Yiming Shao, Kunxue Hong

## Abstract

In our study, we describe a slow progressor CBJC515 from whom we constructed pseudoviruses expressing autologous Env. We surprisingly found all the pseudoviruses were resistant to PGT135. By making site-directed mutations and chimeric Env constructs, we found the early 05 strains escaped from PGT135 by losing the N332 glycan site, while the later 06 and 08 strains may escape with the retention of key epitopes through the change of V1/V4/C2 region or by N398/N611 glycan, which was selected as unique N-glycosylation site of CBJC515 compared with CBJC437 whose viruses were also harboring key epitopes but sensitive to PGT135. These findings provide insights into how HIV-1 can escape from N332-directed broadly neutralizing antibody (bNAb) responses without changing the epitope itself, and these ways may be useful to prolong the exposures of bNAb epitopes and contribute to bNAb development. Furthermore, our chimeric experiments also allowed us to explore the co-evolution and retention of functionality among regions. We confirmed that the V1V2 region has a wide range of effectiveness in interfering with the function of envelope protein and the V3 region can promote protein function recovery and buffer the harmful polymorphisms in the other regions contributing to the Env antigenic diversity. These results may provide some clues for the design of vaccines against HIV-1 strains.

**IMPORTANCE:** Our findings of mechanisms escaping from PGT135 verified the extensive role of long V1 region in mediating escape from V3-bNAbs. In addition, we also found multiple additional ways suggested that extreme variation may be needed by HIV-1 to escape from PGT135 without changing the epitope itself. Although the V3-glycan bNAb responses are among the most promising vaccine targets, as they are commonly elicited during infection, our findings indicated there may be additional difficulties to be taken into account in immunogen design, such as the consideration of other regions and some glycosylation sites affecting the mask of key epitopes, as well as the selection pressure that may be required by other bNAbs. Our chimeric experiment also highlighted the key role of V3 region in contributing to the maintenance of Env diversity by buffering deleterious polymorphisms, which may be helpful for vaccine design.

## Introduction

Inducing antibodies with cross-neutralizing activity is an important indicator for the effectiveness of HIV-1 vaccine and it remains an unsolved challenge, although broadly neutralizing antibodies (bNAbs) developed in 10-30% of HIV-1 infected individuals after several years’ infection, indicating there are no insurmountable barriers for introducing bNAbs in humans by Envelope (Env) (1–5). So far, there is still no immunogen capable of inducing neutralizing antibodies (NAbs) with broad width to tier-2 viruses (6, 7). In 2019, Bricault CA *et al* (8) modified antigens targeting V2-glycan based on the neutralizing signatures of V2-glycan bNAbs, and induced greater breath of tier 2 NAb responses, indicating the potential usage of researching the resistant mechanisms of HIV-1 to bNAbs in vaccine design.

The glycan supersites of the V3 region on HIV-1 Env form vulnerable targets and are exploited by broadly neutralizing monoclonal antibodies such as PGT121, PGT128, and PGT135(9). The N332 glycan residue at the base of the V3 loop has been demonstrated to provide an important supersite of vulnerability for extensive antibody-mediated virus neutralization, and is used to aid the design and development of effective vaccines. In the case of these monoclonal antibodies that target the V3-glycan supersite, the loss of glycan at position 332 is often associated with resistance(10). However, there are still quite a lot of circulating strains that harboring this supersite but still resistant to these antibodies(10, 11). Previous studies have speculated that in the presence of N332 glycosylation sites, the longer length of V1/V2 region was associated with V3-glycan bNAbs resistance (8, 11–14), and van den Kerkhof TL *et al* (11) observed a statistically significant positive correlation between V1 length and neutralization resistance to PGT135. However, these speculations still need to be verified. At the same time, because functionally defined epitopes were distinct from structurally defined epitopes(15), so far, little was known about the determinants conferring neutralization resistance to these viruses. Researching for the mechanisms behind this phenomenon may provide insights for modification of vaccine immunogens and contribute to the design of HIV-1 Env immunogens.

Furthermore, the HIV-1 gp120 protein organization alternates between five constant regions and five variable regions (16) through structural independence, so that the goals of continuous variation and maintenance of function can be ensured. On the one hand, the more accessible variable regions of the immune system can freely undergo extensive sequence diversification to counteract the immune response(17–19); On the other hand, the constant regions are entirely located inside(17, 18), and the more conserved core regions inside provide a scaffold to stabilize the protein structure(20, 21). In the presence of a wide range of co-evolutionary networks, it is especially a challenge to retain functionality. Understanding the coevolution networks is also an important issue for vaccine design(22, 23), which provide insight for the design of antigenic modifications while retaining its functionality.

In our study, pseudoviruses expressing the CBJC515 autologous Env were constructed, and the neutralizing sensitivity of these pseudoviruses to PGT121, VRC01, 12A21, 10E8, 2G12, and PGT135 were tested. Surprisingly we found all of them were resistant to PGT135. Such patient provided us with a unique opportunity to study this phenomenon. Through site-directed mutation and chimeric experiment, we found multiple mechanisms of escape from PGT135, including through long V1 region, changes in V4/C2 region, or through creating 398/611 glycan site, which explored our knowledge especially on how HIV-1 can escape from PGT135 responses through extreme ways without changing the epitope itself. Furthermore, chimeric experiments also allow us to explore the co-evolution and function maintenance among different regions, we verified the V1V2 diversity has broad interference with Env functionality and the role of V3 region in relieving dramatic decrease in functionality induced by V1V2 or C2, which has been rarely studied(24, 25), and these findings will be useful for vaccine design.

## Materials and methods

### Study subject

The donor CBJC515 and CBJC437 described in this study were selected from a Chinese HIV-1 subtype B’ chronically infected cohort, whose plasma exhibit broad cross-neutralizing activity against a panel of 25 viruses since the first sampling year (26). These patients were infected during commercial plasma donation between 1992 and 1995 and were antiretroviral treatment (ART)-naive. The major characteristics of CBJC515 were shown in Table 1, and those of CBJC437 were as previously reported(27). The study was reviewed and approved by the Institutional Review Board of the National Center for AIDS/STD Control and Prevention, Chinese Center for Disease Control and Prevention. The subject provided written informed consent before blood and data collection.

**TABLE 1.**
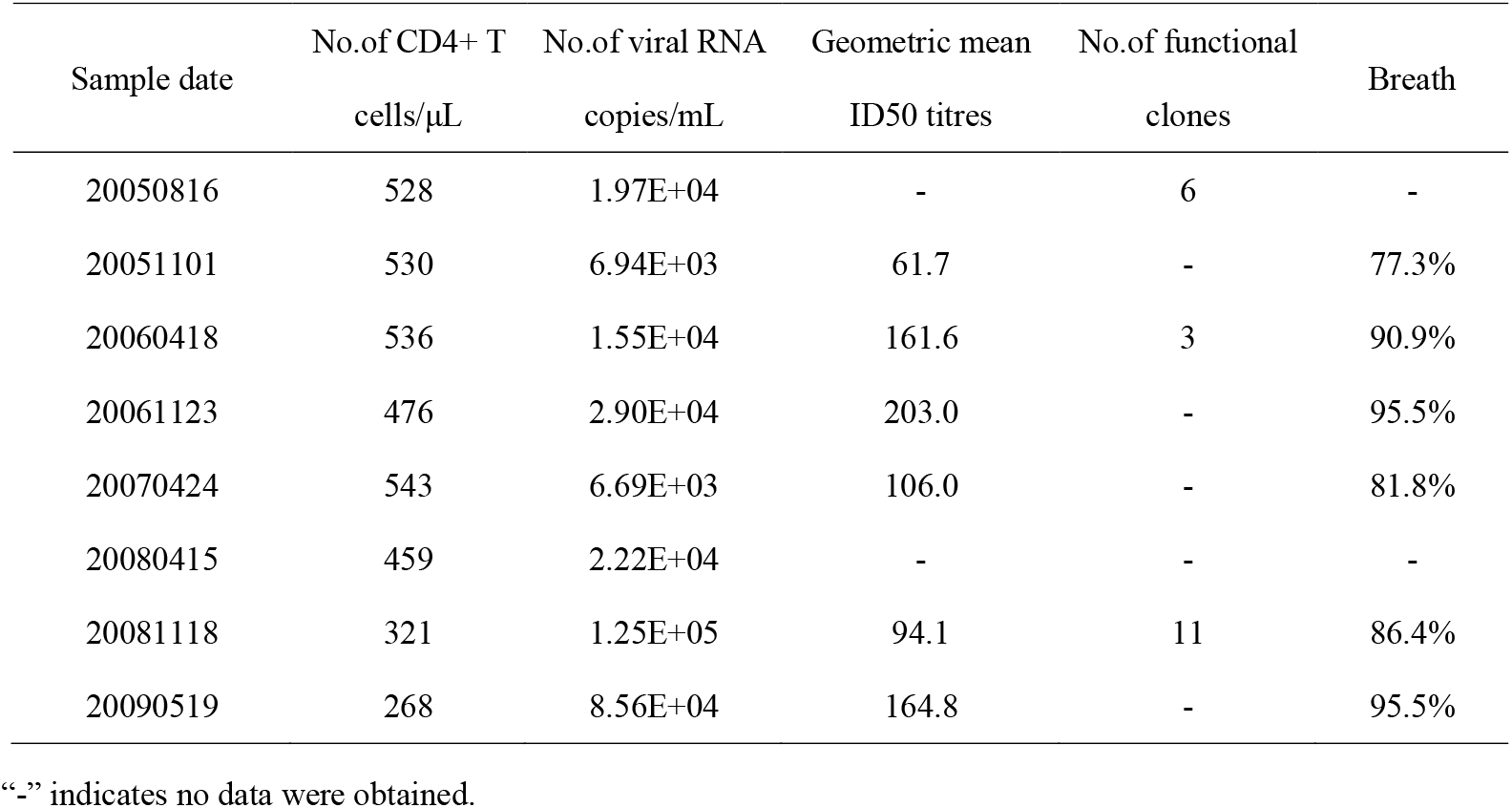
Major characteristics of CBJC515 donor

### Single-genome amplification and envelope clones

Viral RNA was extracted from plasma using QIAamp Viral RNA Mini Kit (Qiagen) and cDNA was immediately synthesized using SuperScript III First-Strand Synthesis System (Invitrogen). Single-genome amplification (SGA) of the full-length gp160 gene was performed as previously described (28). Briefly, the synthesized cDNA was continuously diluted and distributed in replicates of 12 to 16 PCRs in the Thermo Grid 96-well plates to identify a dilution where positive wells accounting for 30% of total PCR reactions. When the SGA criteria of fewer than 30% positive results were met, most of the wells contain amplicons derived from a single cDNA molecule in the appropriate dilution.

The PCR products obtained by SGA were cloned into the commercial directional vector pcDNA3.1D/V5-His-TOPO according to the manufacturer’s instructions (Invitrogen). The correct pcDNA3.1-Env plasmid used to produce wild type pdsudoviruses were selected by sequencing. First, transforming the constructed plasmids into E.coli JM109 competent cells (TaKaRa), and then selecting monoclonal E.coli colonies on ampicillin-resistant LB medium for sequencing. The verified correct E.coli clones were cultivated and then the plasmids were extracted using E.Z.N.A.^®^ Plasmid Mini Kit (Omega) for pseudovirus production. The mutagenesis and chimeric plasmid described below used to produce pdsudoviruses were selected as the same way.

### Sequence alignment and analysis

SGA were sequenced on an ABI 3770 Sequencer (Applied Biosciences). The full-length gp160 gene fragments for each product were assembled and edited using Sequencher 4.1 (Gene Codes, Ann Arbor, MI). All chromatograms were inspected for sites of mixed bases (double peaks), and any sequence with evidence of double peaks was excluded from further analysis.

The Env nucleotide sequences were aligned by Gene Cutter (https://www.hiv.lanl.gov/content/sequence/GENE_CUTTER/cutter.html) using HIV-1 HXB2 as reference sequence and the amino acid sequences were deduced by Env nucleotide sequences. Potential N-linked glycosylation sites (PNGS) were identified using N-Glycosite (http://www.hiv.lanl.gov/content/sequence/GLYCOSITE/glycosite.html) at the Los Alamos HIV database website; The consensus sequence of CBJC515 and CBJC437 were analyzed by online analysis tool “Consensus Maker” (https://www.hiv.lanl.gov/content/sequence/CONSENSUS/consensus.html); LALIGN tool (http://www.ch.embnet.org/software/LALIGN_form.html) was used to calculate sequence identity and similarity between the 2005 Env consensus sequence and each 2005 functional clones; Chinese subtype B database set (named “B-Database”) containing a total of 168 sequences were selected from Los Alamos HIV database (https://www.hiv.lanl.gov/components/sequence/HIV/search/search.html) according to the keywords “china”, “subtype B”, “intact gp120 sequences” and “one sequence/patient”.

### Generation of site-directed mutagenesis and chimeric clones

Site-direced mutagenesis was performed by standard PCR procedure. Approximately 50 ng plasmid DNA template, 1 μL of 1 μM primer F (forward), R (reverse), were added into the PCR mixture containing 4 μL dNTP mixture (2.5 mM), 25 μL 2×PrimeSTAR GC buffer (TaKaRa), 0.5 μL PrimeSTARWHS DNA polymerase (2.5 U/μL) (TaKaRa) and ddH_2_O, with a total volume of 50 μL. The primer F was designed according to the nucleic acid sequence of 5 amino acid positions before and after the mutation site, and the primer R was reverse and complement to the primer F. Cycling conditions for this PCR are 98°C for 1 min, followed by 30 cycles of 98°C for 10 s, 56°C for 20 s (the annealing temperature can be adjusted according to different primers), and 72°C for 6 min, with a final extension at 72°C for 10 min. The entire Env gene of each mutant was sequenced to confirm mutation.

Chimeric clones were conducted by GeneArt^®^ Seamless Cloning and Assembly Kit (Invitrogen), which can assemble several linearized DNA fragments into one DNA sequence. The inserted nucleotide fragments were synthesized (Sangon Biotech). The backbone of Env plasmid to be replaced were linearly amplified by PCR, and the PCR products were verified on a 0.8% agarose gel and then subjected to gel extraction using QIAquick Gel Extraction Kit (Qiagen). The two ends of the linear amplification primers were designed to overlap with synthesized insertion protein sequence with 15bp bases respectively, so the two parts were able to be seamlessly linked through the assembly kit (Invitrogen). The integrity of the chimeric plasmids was confirmed by sequencing and restriction analysis as described above. Finally, the boundaries of regions changed after chimeric were as follows: 131-135 for V1 region, 158-196 for V2 region, 131-196 for V1V2 region, 197-295 for C2 region, 296-331 for V3 region, 385-418 for V4 region, and 131-331 for V1-V3 region. The numbers refer to the amino acids position of HXB2 gp120 protein.

### Pseudovirus preparation, and titration for neutralization assays

Pseudoviruses were prepared, titrated as previously described(27). Briefly, 293T cells were cotransfected with pcDNA3.1-Env clone and Env-deficient HIV-1 backbone vector (pSG3△Env) using PEI transfection reagent (PolyScience). Pseudovirus-containing supernatant was harvested 48h post-transfection, and then filtered (0.45μm pore size) and stored at −80°C. The 50% tissue culture infectious dose (TCID50) of a single-thawed aliquot of each pseudovirus batch was determined in TZM-bl cells.

### Neutralization assay

Neutralization was measured as a reduction in Luc reporter gene expression after a single round of virus infection in TZM-bl cells as described previously (26). Briefly, 50 μL of pseudotyped viruses normalized to 4000 TCID50/mL was incubated with 100 μl serial threefold dilutions of broadly neutralizing monoclonal antibodies (bN-mAbs) in duplicate for 1 h at 37°C in 96-well flat-bottom culture plates. The virus-antibody mixture was then used to infect 10,000 TZM-bl cells in the presence of 30 μg/mL DEAE-dextran. One set of the control wells received cells only, while the other set received pseudovirus plus cells. Infection levels were determined after 48h by measuring the luciferase activities of cell lysates. After 48 hours of incubation, 150 μl of the culture was removed, and 100 μl of Ultra-High Sensitivity Luminescence Reporter Gene Assay System (PerkinElmer) was added and incubated for 2 minutes. luciferase activities of 150 μl lysate transferred from each well to a 96-well black solid plate were measured by a luminometer (PerkinElmer). The 50% inhibitory dose (ID50) was defined as either the plasma dilution or sample concentration at which relative luminescence units (RLU) were reduced 50% compared to virus control wells.

### Neutralizing antibodies used in the study

bN-mAbs PGT135, PGT121, 2G12, 12A21, 10E8, and VRC01 were kindly received from NIH AIDS Research and Reference Reagent Program.

### Data availability

Env sequences from CBJC515 and CBJC437 donors have been deposited into GenBank and the accession numbers were shown in Table 2 and Supplementary Table 1.

**TABLE 2.**
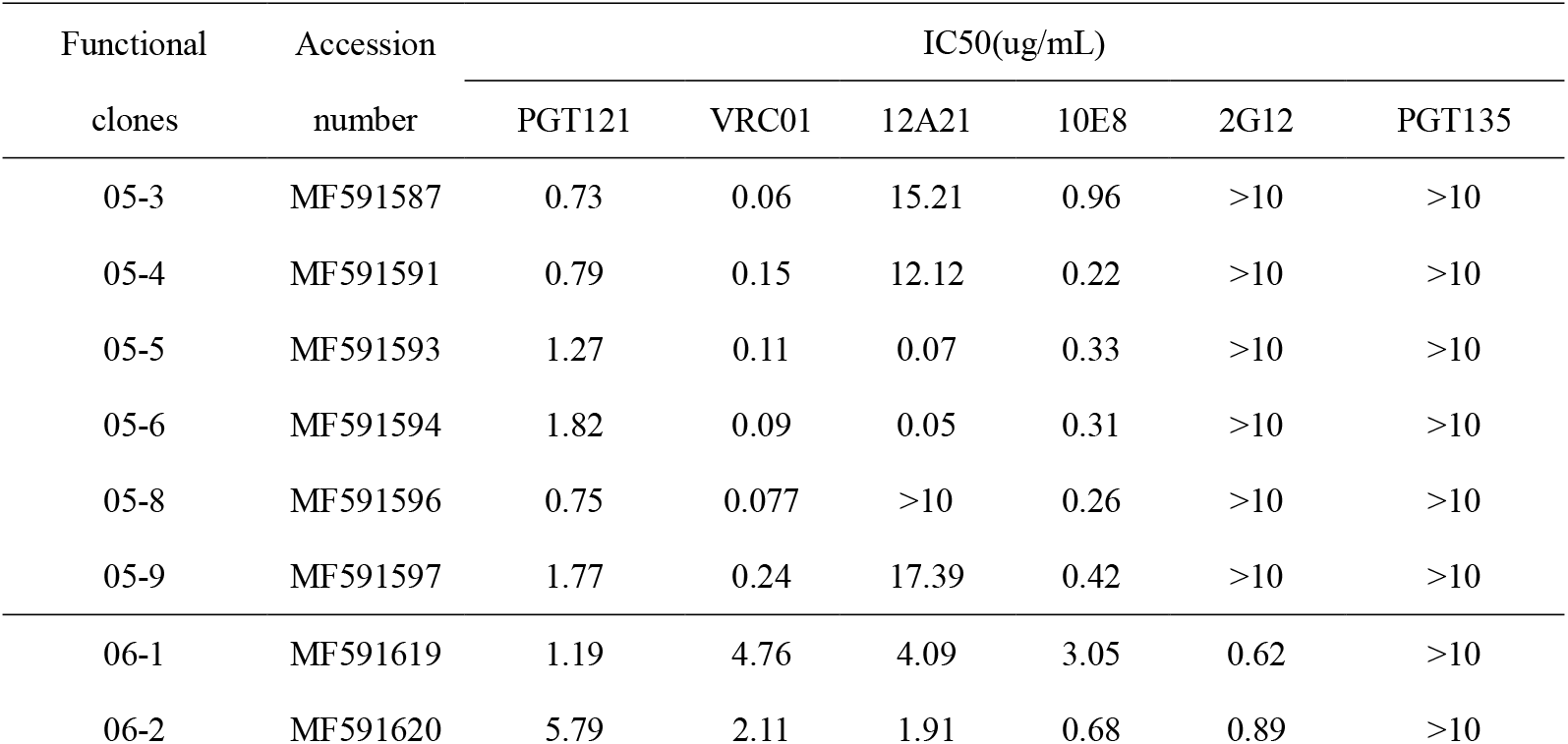
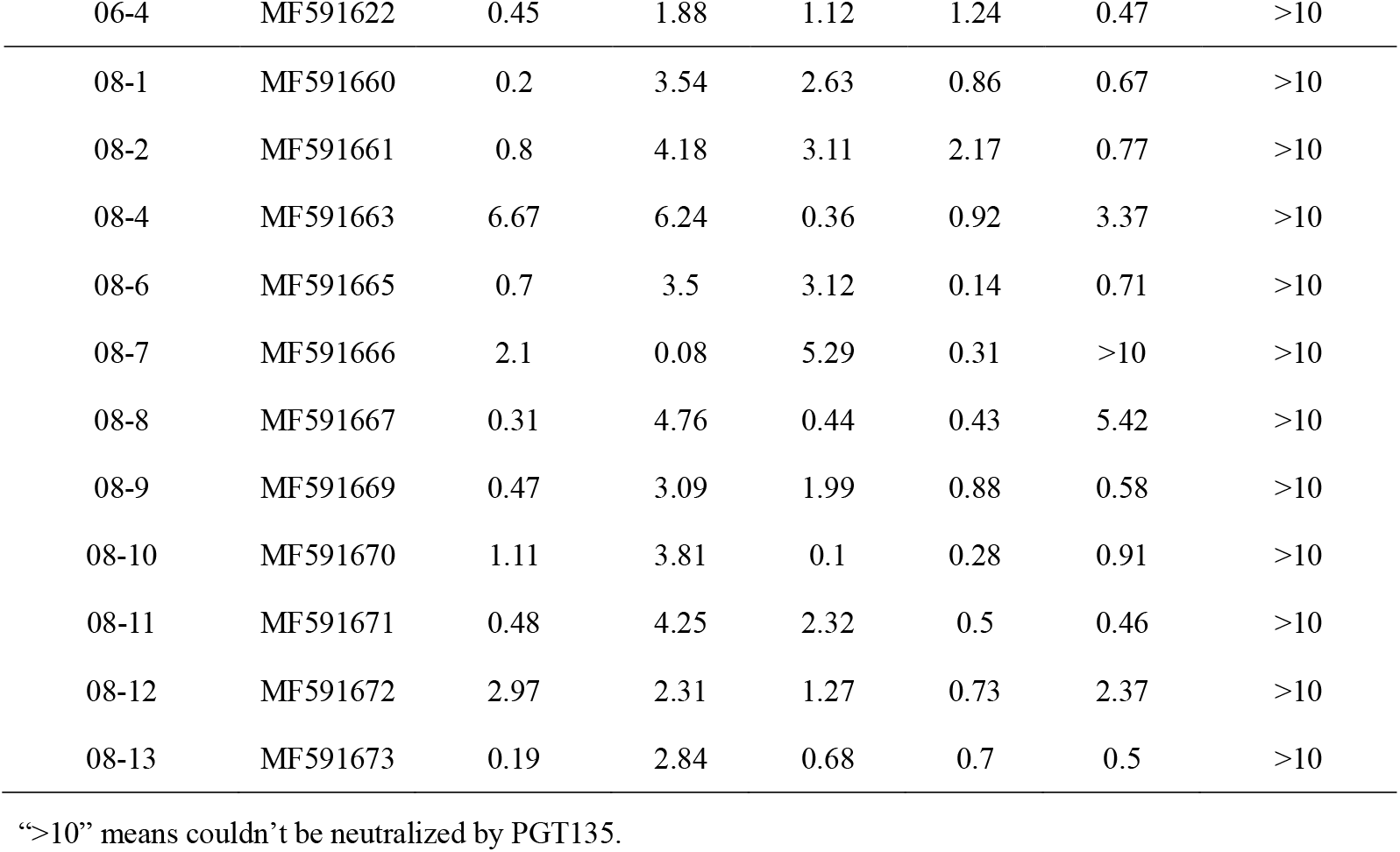
Neutralization sensitivity of CBJC515 pseudoviruses to representative broadly neutralizing monoclonal antibodies (bN-mAbs)

## Results

### Sensitivity to neutralization by broadly neutralizing monoclonal antibodies (bN-mAbs)

6, 3, 11 pseudoviruses were successfully constructed from CBJC515 plasma samples at 20050816, 20060418, and 20081118 time points, named 05/06/08 isolates respectively, and we examined the sensitivity of these isolates to prototypic bNAbs PGT121, VRC01, 12A21, 10E8, 2G12 and PGT135 (Table 2). Almost all 20 clones were sensitive to the CD4 binding site (CD4bs) specific bN-mAb VRC01, 12A21, and membrane proximal external domain (MPER) specific bN-mAb 10E8. As to the bN-mAbs recognizing N332-supersite (9, 29), all clones were resistant to PGT135 but sensitive to PGT121, while all 05 strains were resistant to 2G12.

### Lack of 332 glycosylation site leads to resistance to PGT135 in 05 isolates

Since all clones from CBJC515 were resistant to the V3-glycan bNAb PGT135, this aroused our great interest. PGT135 interacts with glycans at Asn332, Asn386 and Asn392, using long CDR loops H1 and H3 to penetrate the glycan shield to access the gp120 protein surface(9). We first analyzed the 332 position of these strains. As shown in Table 3, the N332 glycan site was generally missing in 05 Env clones, but all 06/08 clones contained this supersite. In 05 isolates, strain 05-8 lacks 332 N-glycan site due to the presence of aspartic acid at this position (referring to the relative position on the HXB2 strain, the same below), while others possess N334 glycan site, which also leads to lack of 332 N-glycan site. We performed D332N or N334S mutations on 05 isolates, and all of them became sensitive to PGT135 and 2G12 (Table 4). In addition, we checked other epitopes reported in literatures (8, 9, 30) that associated with PGT135 neutralizing sensitivity(Table 3), and performed R/K389Q, T409E or Y330H mutation to the corresponding strains, but no changes were found.

**TABLE 3.**
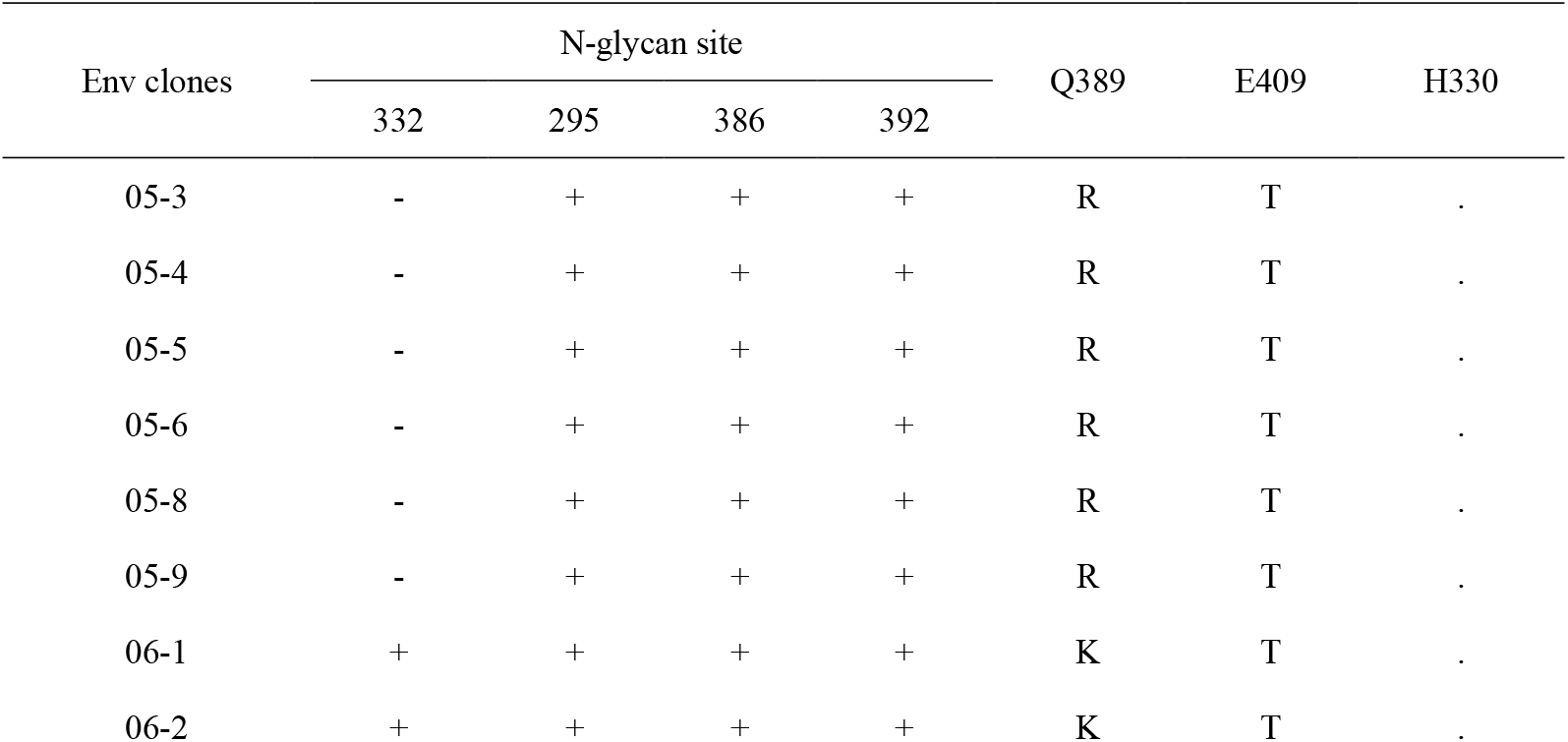
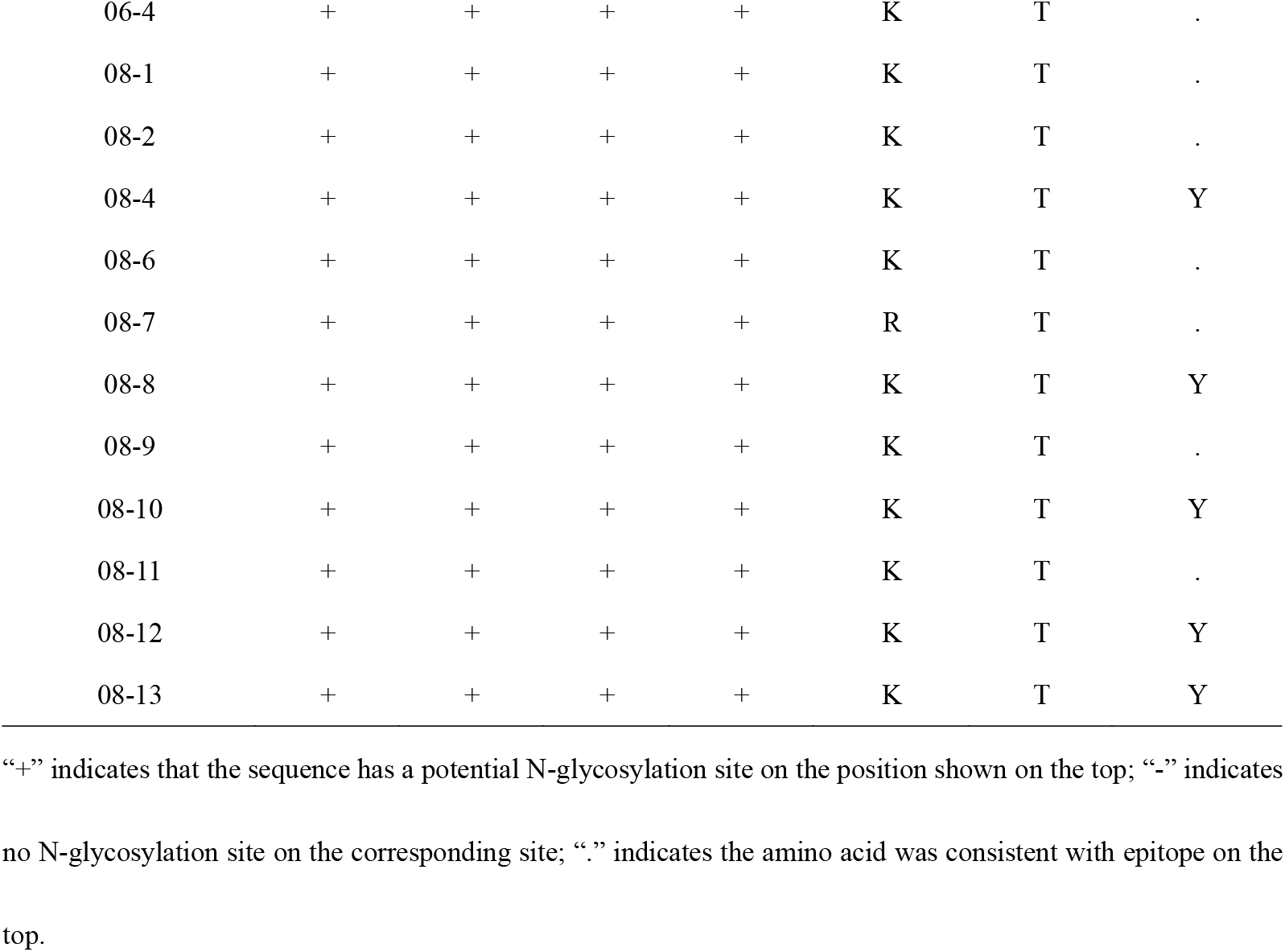
The condition of epitopes for PGT135 in 2005, 2006 and 2008 functional Env clones

**TABLE 4.**
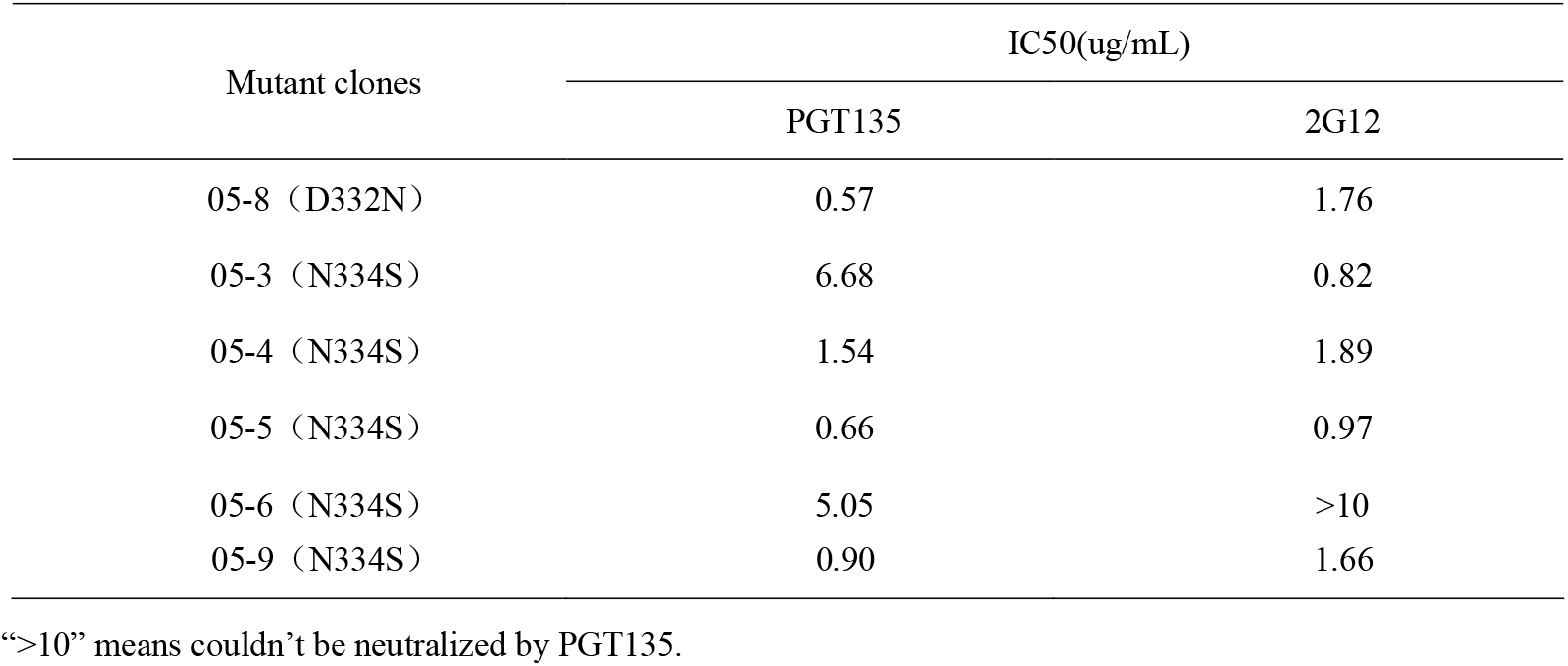
The neutralizing sensitivity of mutations to PGT135

In our study, PGT135 and 2G12 were completely unable to neutralize the 05 pseudoviruses containing N334 glycan site, while PGT121 was able to neutralize all of them. Previous studies have shown that some N332-dependent bNAbs can recognize promiscuous glycans with different ability, and some can neutralize strains transferred glycosylation site to N334(9, 10). Our observations were consistent with it.

### Chimeric experiment examines effect of different regions on neutralizing sensitivity of 06/08 isolates

We further explored the determinants conferring neutralization resistance to 06/08 strains with key epitopes (Table 3). Sequence analysis showed that 05 isolates had shorter V1 region of 22 amino acids (aa) with higher similarity, while the 06/08 strains expanded V1 region pronouncedly (22-42 aa long) and was much longer than HIV-1 subtype B consensus sequence (25 aa) (13) (Fig. 2). By replacing the V1 region (position 131-157) of 06/08 Env clones to 05-8 (D332N), which was selected as the most representative strains of 05 clones through sequence harmony (SH) method compared with 05 Env consensus sequence (Table 5), we assessed whether longer V1 loop was associated with neutralizing resistance to PGT135 (11, 13).

**FIG 1.**
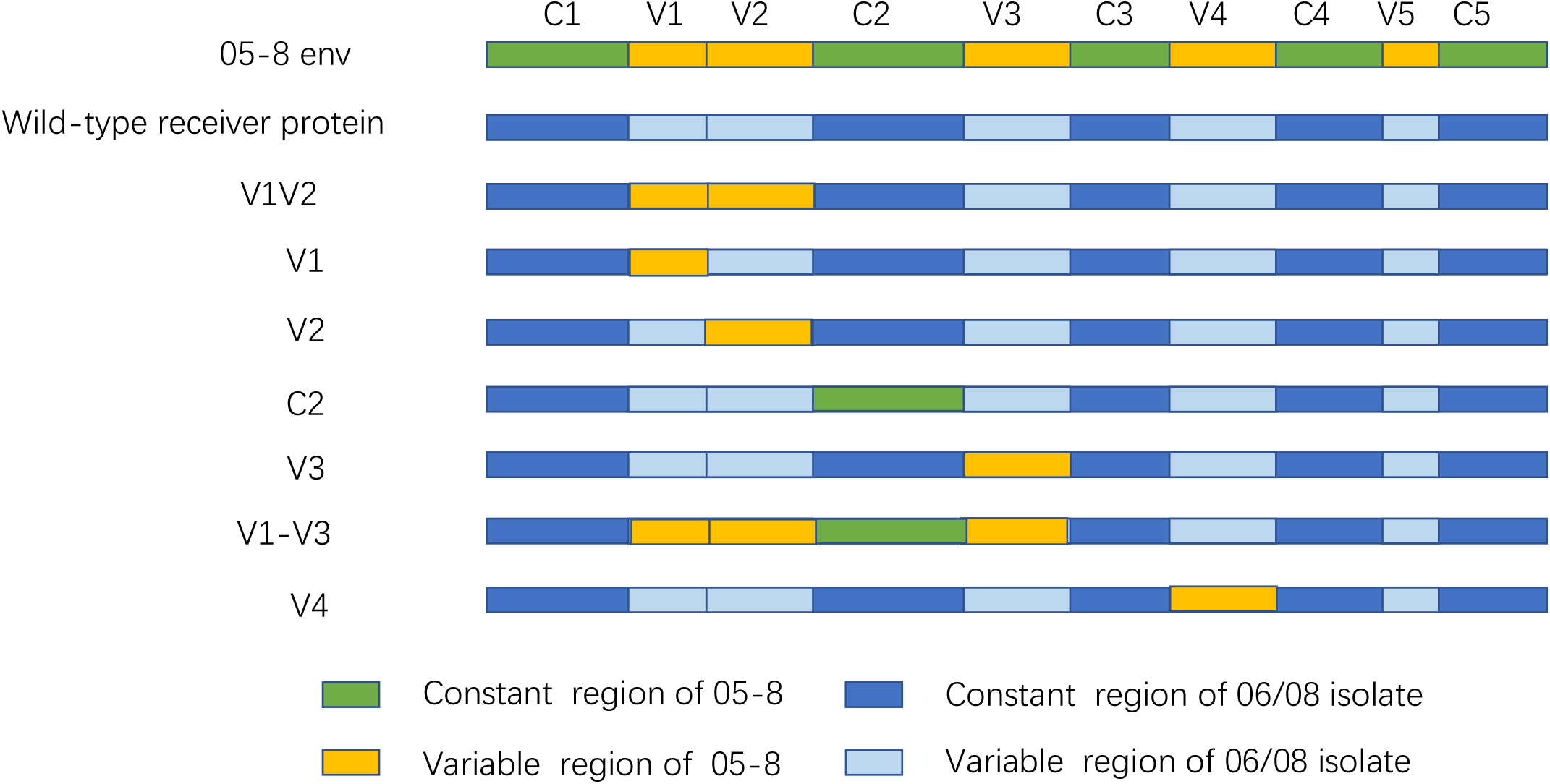
Schematic representation of the chimeric HIV-1 envelopes used in this study. The C1-C5 structure of the 05-8 envelope gene is given at the top, with the constant and variable regions of the gp120 region in green and yellow, respectively. In the 06/08 wildtype receiver protein, the backbone of constant and variable region is shown in dark blue and light blue, respectively.

**FIG 2.**
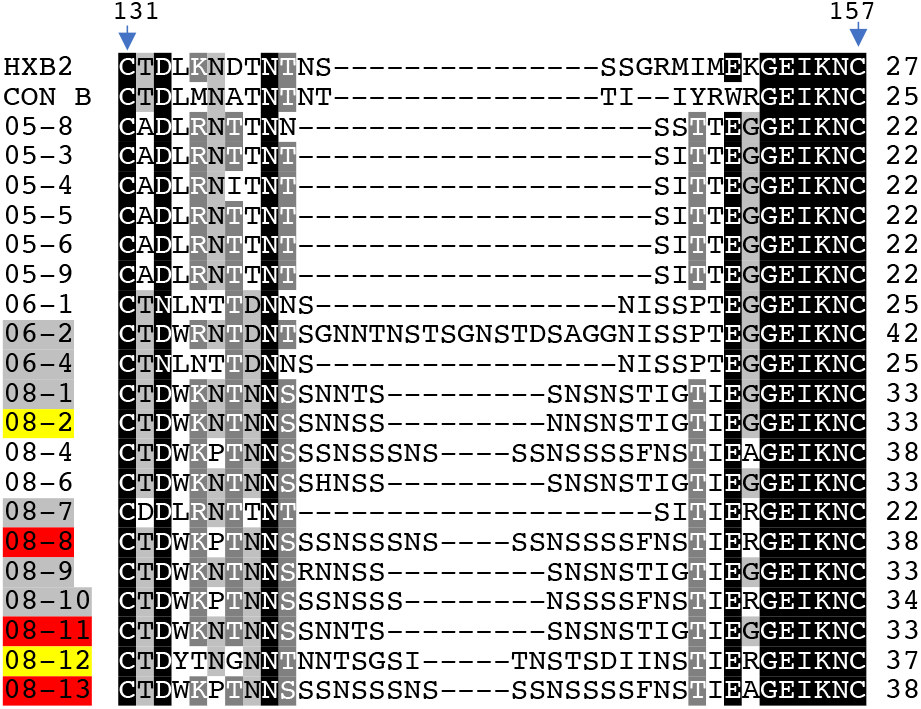
The alignment of V1 region. CON B representing the HIV-1 subtype B consensus sequence was from reference(13). Sequence position corresponding to reference strain HXB2 is 131-157. The nucleic acid with gray, dark gray, and black background represents common amino acid sequences. Gaps in the amino acid sequence are shown with a dash (-). The rightmost number shows the length of V1 region. The names of sensitive chimeras are in red background, resistant chimeras are in yellow background, and non-functional chimeras are in gray background.

**TABLE 5.**
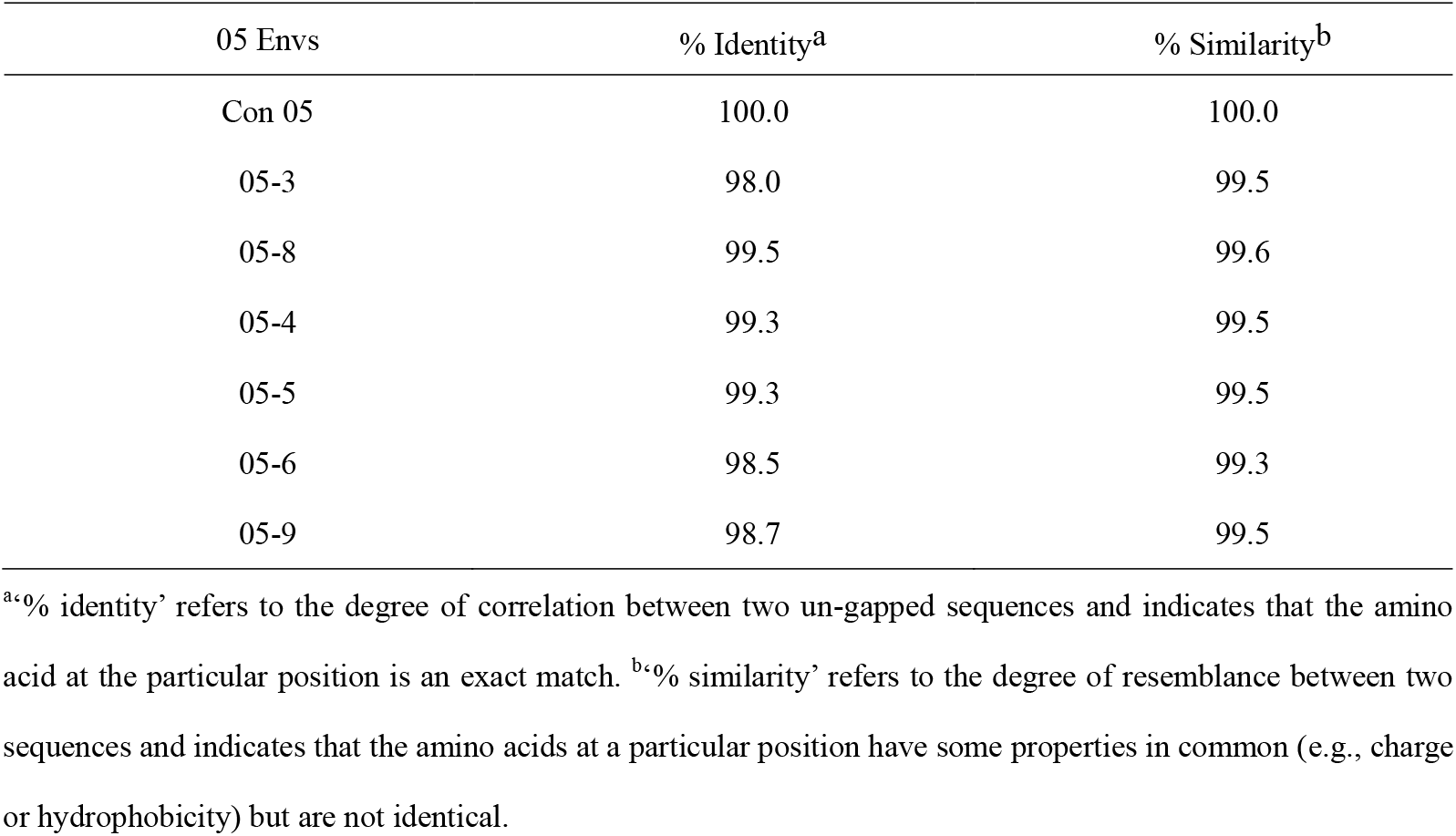
Similarity of amino acid sequence of consensus 05 funcional Env with other 05 functional Envs

11 V1 chimeras were successfully constructed (Fig. 2 and Table 6). Among them, 6 chimeras were not able to produce infectious clones (the wild type V1 length varied from 22 to 42 amino acids), 2 chimeric strains 08-2 (05-8 V1), 08-12 (05-8 V1) were still resistant to PGT135 (the original V1 length was 33, 37 amino acids respectively), and 3 chimeric strains, 08-8 (05-8 V1), 08-11 (05-8 V1) and 08-13 (05-8 V1), became sensitive to PGT135 with the IC50 of 1.19 ug/ml, 5.8 ug/ml, and 8.3 ug/ml.

**TABLE 6.**
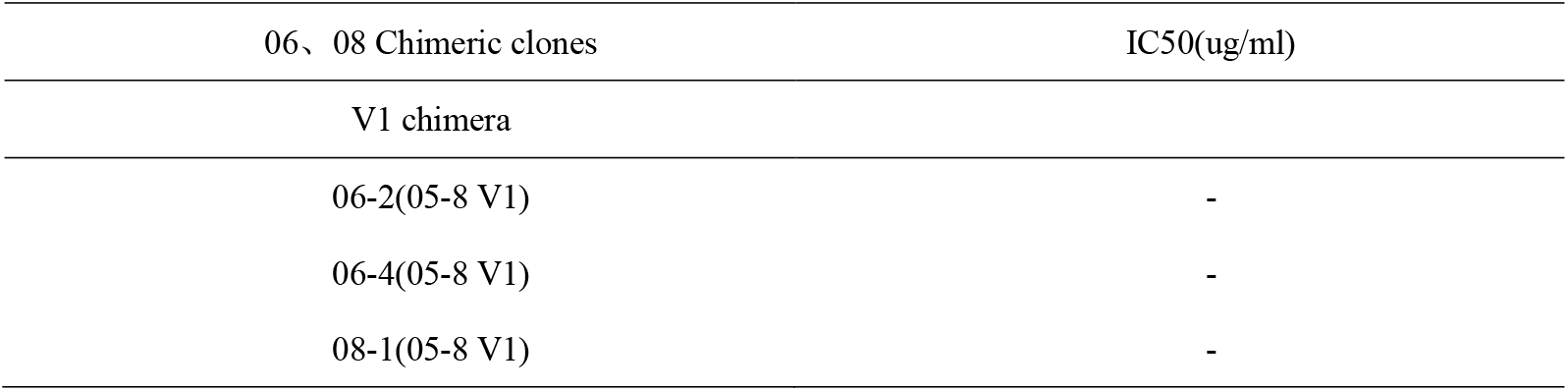
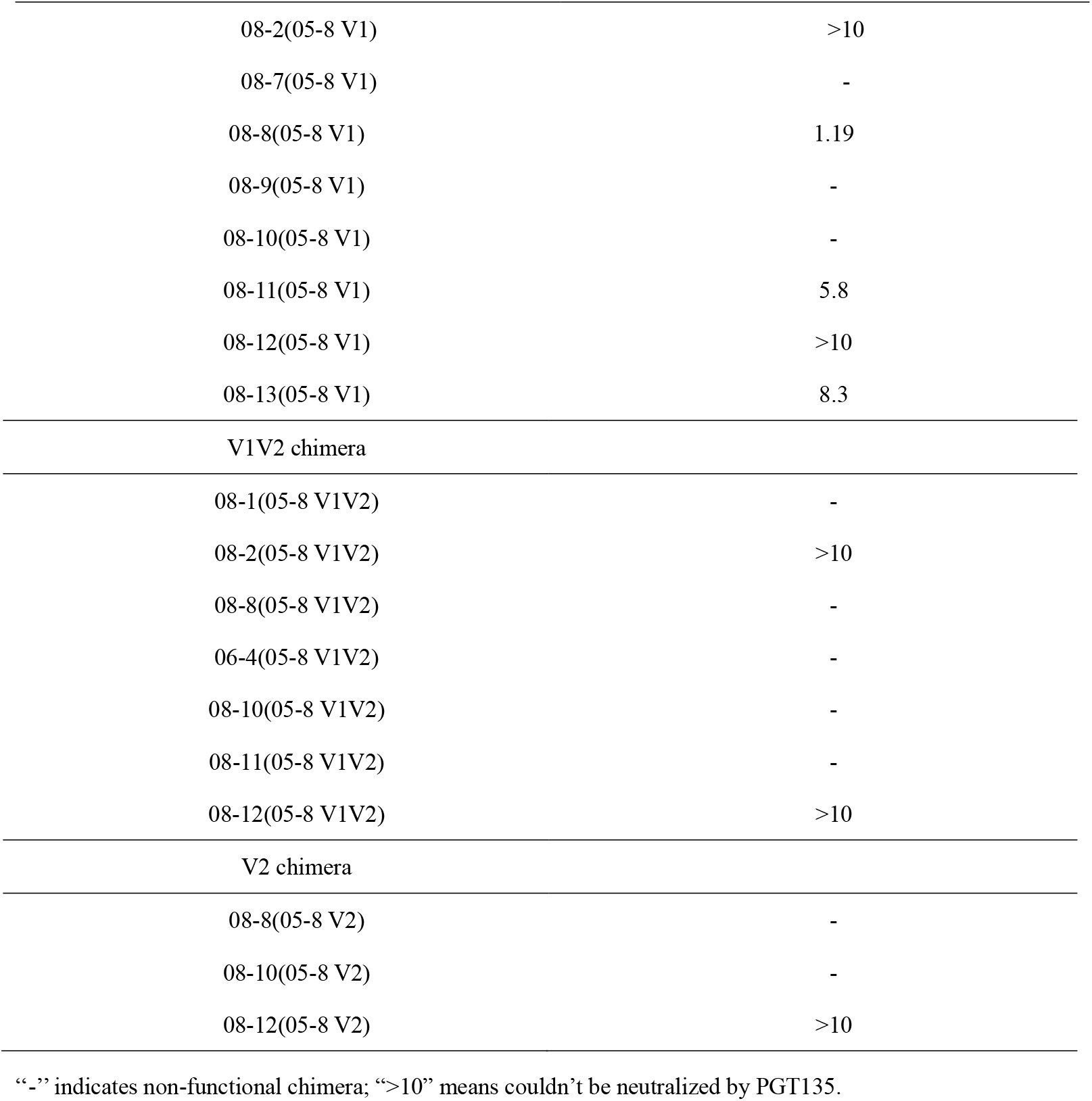
Neutralizing sensitivity of V1/V1V2/V2 chimeras to PGT135

Since the V1V2 region is a global regulator of neutralizing sensitivity(31–34), previous studies also shown that the neutralizing ability of some V3-glycan bNAbs were associated with the length of V1V2 region in the presence of N332 glycan site, and the longer the virus in the V1+V2 region, the more difficult it was to be neutralized (8). Therefore, we also construct 7 V1V2 chimeras by replacing the V1V2 region of 06/08 strains with that of 05-8 (Table 6). Our neutralizing experiment showed 5 of them lose functionality, and only two chimeras 08-2 (05-8 V1V2) and 08-12 (05-8 V1V2) were functional but still resistant. Furthermore, we constructed 3 V2 chimeric clones (Table 6), and there was only 1 chimera was functional and remaining resistant. So, it seems like, among the V1V2 region, V1 loop was the most important part to confer resistance to these strains.

Based on the above results, we further examined whether changes in other regions except for V1 would affect its sensitivity to PGT135 as the virus evolving at subsequent 06/08 time points. Chimeric envelopes were generated by replacing the C2, V3, V1-V3 or V4 region of 06/08 isolates with the corresponding region of 05-8 (D332N) (Table 7 and Fig. 1). Neutralization experiment showed all the 6 V3 chimeras did not change sensitivity to PGT135 (Fig. 3). In the 4 C2 chimeras, 3 were functional and 1 was sensitive. 10/12 V1-V3 chimeras were functional with 3 of them became sensitive, while 2/3 V4 chimeras were functional and all the 2 chimeras were sensitive.

**FIG 3.**
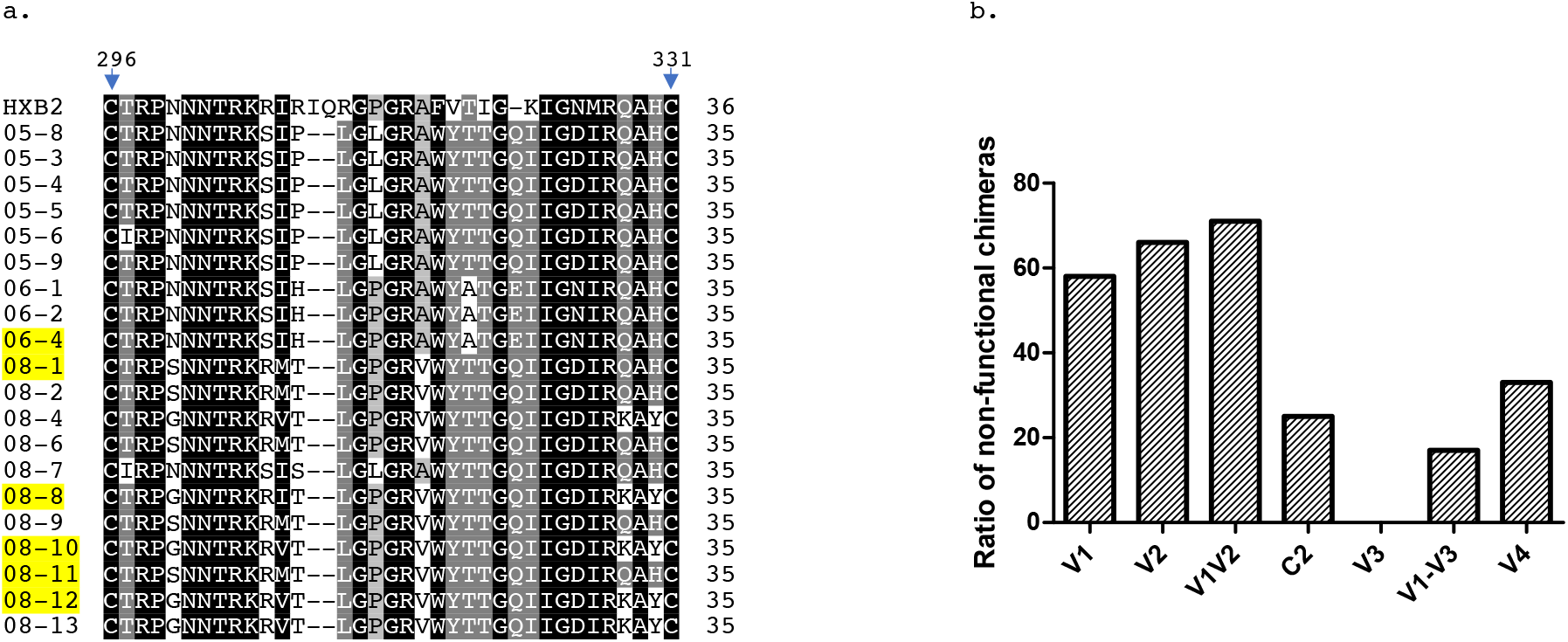
V3 alignment (a) and ratio of non-functional chimeras when chimeric with V1, V2, V1V2, C2, V3, V1-V3, and V4 region (b). The nucleic acid with Gray, dark gray, and black background represent common amino acid sequences. Resistant chimeras are in yellow background.

**TABLE 7.**
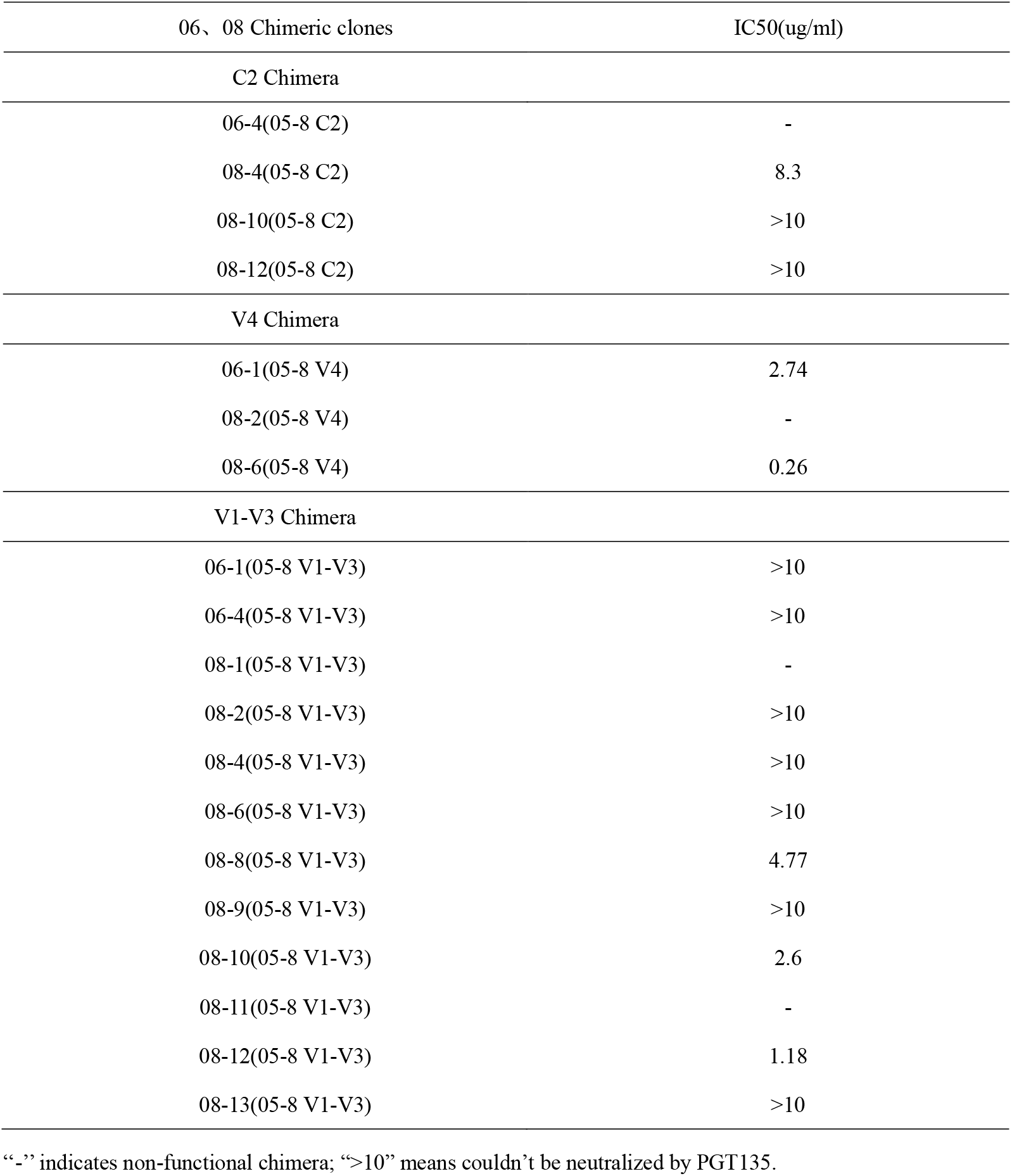
Neutralizing sensitivity of C2/V4/V1-V3 chimeras to PGT135

### N611A and N398A mutation may have Impact on Neutralizing sensitivity to PGT135

To further explore the resistant mechanisms in 06/08 isolates, we selected another Chinese subtype B’ slow progressor CBJC437, whose isolates were also containing key epitopes such as N332/N386/N392/N295/H330 and were sensitive to PGT135 (Table S1). According to the consensus sequence alignment, we selected the different sites of the two samples in V3/V4 region (Fig. 4), which was most likely to influence the neutralizing sensitivity(9), to perform mutations on 3-4 strains of CBJC515 changing the amino acids to the corresponding signature of CBJC437 isolates. However, no mutations were found to have effect on the neutralization. Furthermore, we also compared all the N-glycosylation sites among the two samples, and selected the unique N-glycan site of each sample (since the V1 region has changed a lot, we didn’t consider the unique glycosylation sites in the V1 region) (Fig. 4). We only found the N398A mutation in 06-4/08-13 and N611A mutation in 08-9 restored sensitivity to PGT135 with the IC50 of 7.23 ug/ml, 6.91 ug/ml and 2.64 ug/ml respectively, in a strain-specific way (Table S2-S3).

**FIG 4.**
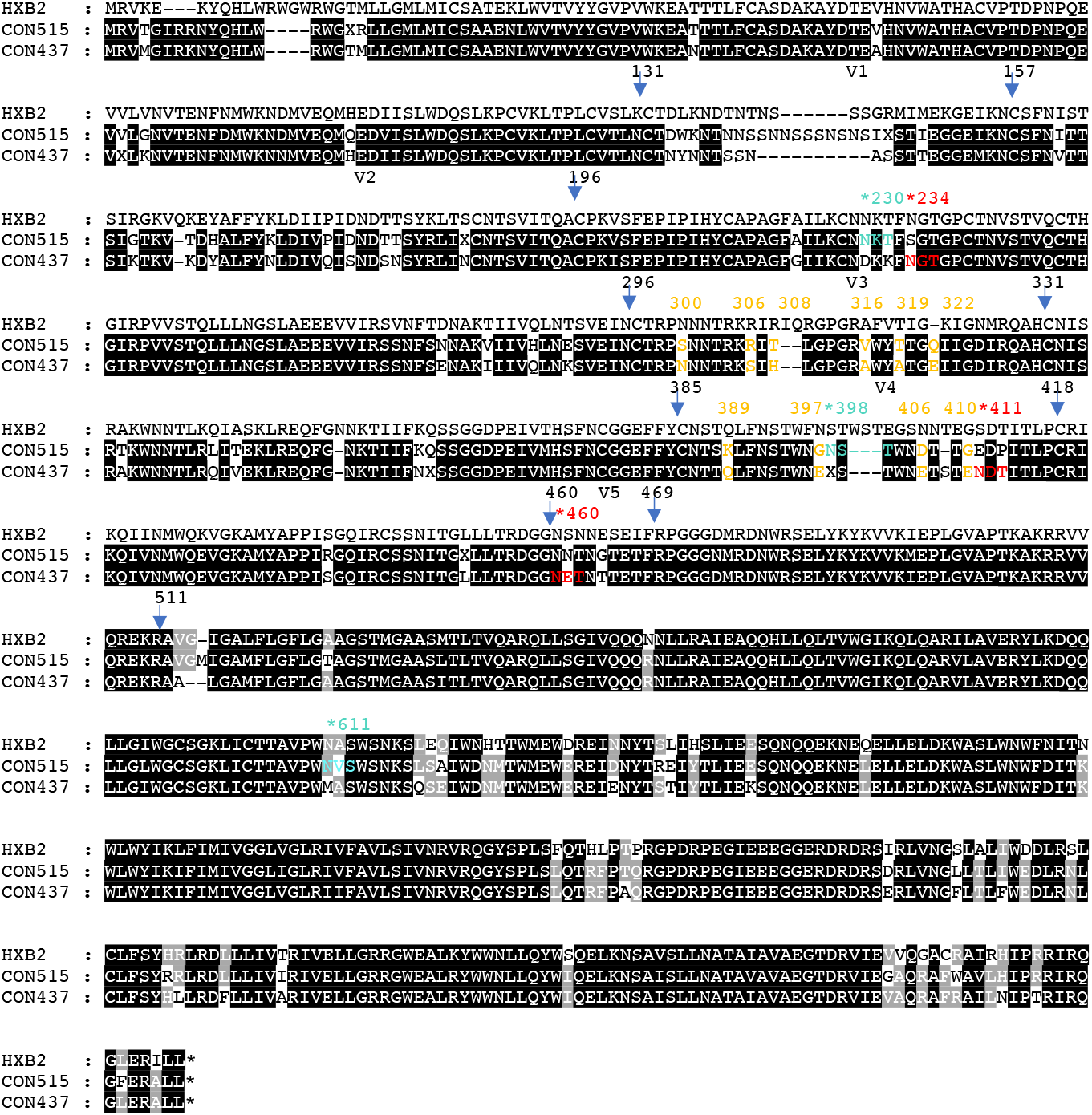
Alignment of the consensus sequence between CBJC515 and CBJC437. Numbers with Asterisks represent potential glycosylation sites; Blue fonts and red fonts represent CBJC515-specific and CBJC437-specific glycosylation sites (excluding V1 region) respectively by N-Glycosite analysis; Yellow fonts represent different non-glycosylation sites among V3/V4 regions between CBJC515 and CBJC437 consensus sequence. The arrows and the above numbers represent the boundaries of each variable and constant region.

### Statistical analysis of key glycosylation sites of PGT135 in Chinese HIV-1 subtype B strains

According to previous reports, the neutralization breadth of PGT135 was about 33%, which was comparable to the CD4bs-class antibody b12, but lower than other N332-dependent antibodies such as PGT121 and PGT128 (29). This lower neutralization width was reported mainly due to the limited prevalence of key contact residues, such as Asn332, Asn392 and His330, in circulating strains(9).

We downloaded 168 sequences of Chinese subtype B strains from Los Alamos HIV database (named B-Database), and made statistics on the key neutralizing epitopes of PGT135. H330, N332 and N392, which are necessary for PGT135 neutralization, were present in 76% (127/168) sequences of B-Database. As PGT135 can recognize N295 and N386 glycans in a strain-dependent manner(9), we also calculate the prevalence of H330, N332, N392 and N295 or N386, and the ratio was 66% (111/168) and 67% (112/168) respectively. At the same time, all the four N-glycan site, N332 N392 N295 and N386, appeared in 60% (100/168) of the sequences, and the ratio of single glycosylation site at position 332 accounted for 93.5% (157/168). The prevalence of these key sites was significantly higher than the 33% neutralization width of PGT135. It can be speculated that a large part of the Chinese HIV-1 subtype B strains containing key epitopes of PGT135 may able to escape from it. Therefore, it will be very meaningful to explore the resistant mechanisms of these strains.

## Discussion

The generation of bNAb responses through vaccination is still an essential question in HIV research. Knowing the escaping mechanisms in broad cross-neutralizing samples may yield useful information for vaccine design (14, 35–37). In our research, escape from PGT135 was mediated via multiple pathways. Except for the absence of N332-supersite, the resistant strains harboring all key epitopes may escape through changing V1/V4/C2 region, or through N398/N611glycan. Previous study speculated strains may escape from PGT135 through longer V1 region(11), and in our research, we verified it can indeed restore the sensitivity in some strains, while the V1V2 or single V2 replacement cannot. Together with previous study about PGT121 and PGT128(10, 12, 38), our finding highlighted the broad effect of long V1 region on neutralizing activity to V3-glycan bNAbs which may through blocking access to the V3-glycan supersite(14). We also found V4 and C2 region may affect its neutralization. Through our large amount of site-directed mutations, we found it very difficult to restore neutralizing sensitivity by changing single amino acid. Our observations thus suggested that extreme variations may be needed to escape from PGT135 in strains with N332-supersite(13).

In the CBJC515 donor, over the course of infection, the locations of PNGS were altered. Strains evolved from 2005 strains without N332 glycosylation site to the isolates with 332 N-glycan gradually fixing in 2006/2008 timepoint. The usual mode of escape from the V3-carbohydrate class of potent broadly neutralizing antibodies is via mutation of the N332 glycan(10), so it can be speculated that samples from time points prior to 2005 may contain V3-glycan bNAbs such as PGT135-like. After the year 2005, We speculated that the retention of the N332 epitope may be due to selection pressure from strain specific NAbs or other antibody lineages (14, 35) as time went on. In order to escaping from them, strains produced N332-glycosolation site and adopted these unusual escaping routes. The neutralization width of CBJC515 fluctuated continuously from 2005 to 2009 timepoint (Table 1) and previously study which suggesting different lineages of antibody co-exist in this sample(39) supporting our prediction. In natural infection, bNAbs usually take several years to evolve, suggesting a requirement for prolonged antigen exposure (36, 40, 41). In CBJC515, the retention of N332 site through extreme escape routes may help to prolong the exposure of bNAb epitopes, which may assist its maturation as mentioned in previous reports (14, 35–37), and it was supported by our results of incomplete viral escape in contemporaneous neutralization (data not shown). The explanation and speculation of the unusual escaping mode in CBJC515 donor may need to be further explored, such as by plasma epitope mapping and B-cell sequencing.

Genetic diversity and coevolution are essential for HIV-1 to get rid of the host immune response. Extensive coevolution of structurally and functionally related parts of the protein is the solution for the virus to conciliate genetic diversity and maintain functionality. For these aspects, there have been few reports in the past(24, 25), and our chimeric experiment provided materials for the investigation.

As the outermost part of the viral particle, gp120 is the virus component most vulnerable to immune stress, and therefore, may be the most important protein for genetic diversity. Although V1V2 region might preserve the functionality of the protein with only minor structural constraints while they have evolved to maximize their possibility of sequence diversification, the replacement of variable region abolishes Env functionality in 5/7 V1V2 chimeras, 6/11 V1 chimeras and 2/3 V2 chimeras (Table 6 and Fig. 3b), which indicated the broader validity of V1V2 region in interference with the functionality of envelope protein (25). In addition, there were 1/3, 1/4 non-functional clones in V4, C2 chimeras, respectively (Table 7 and Fig. 3b). The non-functional clones after the replacement of these regions reflect the important significance of co-evolution among each region of the strain for function maintenance.

As it has been reported that the replacement of V3 usually has less impact on Env function(25), we did not find any non-functional clones through chimeric V3 region (Fig. 3a). The higher conservation level of V3 region may reflect the requirement to maintain the optimal conformation required for the co-receptor binding site and explain why the replacement of this domain is more tolerant than other regions(42). When we replace the V1-V3 region (including V1, V2, C2, V3), there were only 2/12 chimeras showed non-functionality (Fig. 3b). This extremely low probability of non-functionality compared with the single exchange of V1/V2/C2 region, in addition to reflecting the coevolution of V1-V3 region, it also reflects the role of V3 in promoting function recovery. Taking the 06-4 strain as an example, chimeric with V1V2 and C2 region all abolished its functionality, while V1-V3 chimeras restored. A recent study support our finding of the role of V3 region in buffering deleterious effect of polymorphisms and increasing genetic robustness (24), and this study also pointed out that the defect of C2 chimera may be blocked after recognizing the co-receptor CCR5 due to interference with the subsequent conformation of membrane fusion changes.

Finally, we deepened the understanding of the pathways that may affect the accessibility of PGT135 to its epitopes and highlighted the complexity and extremely ways of resistant mechanisms to PGT135 under the condition of harboring key epitopes which verified and supplemented the previous results. We speculated that these unusual ways may play an auxiliary role in the development of bNAb and may be related to the maintenance of long-term highly broad cross-neutralizing activity, although it needs further exploration. The chimeric experiment also verified the importance of coevolution of structurally and functionally related parts of Env for the virus to conciliate genetic diversity and maintain functionality, especially the role of V3 region in the retention of functionality. Altogether, our findings may be helpful for the design of antigenic modifications while retaining its functionality.

## ACKNOWLEDGMENTS

This work was supported by the grant from the National Natural Science Foundation of China (81172809, 31411130194), the National Major Project for Infectious Disease Control and Prevention (2016ZX10001-008, 2018ZX10731-101), the SKLID key project (2019SKLID602) and International Cooperation Grant from the Ministry of Science and Technology of China (2016YFE0107600). We are grateful to the CAVD HIV Specimen Cryorepository (HSC) for their contribution of HIV-1 Env-pseudotyped viruses for this study.

## AUTHOR CONTRIBUTION STATEMENT

SS, LM, YS and KH conceived and designed the study. SS, SZ, YH, YL, LR and YLH performed the experiment. SS, YL, XH, YR and KH analyzed the data and edited the manuscript. All authors have read and approved the final manuscript.

